# WarpDemuX-custom - Multiplexed sequence-specific direct RNA sequencing of ribosomal RNA and beyond

**DOI:** 10.64898/2026.10.09.757944

**Authors:** Isabel S. Naarmann-de Vries, Wiep van der Toorn, Swastik Mishra, Maximilian F. Blank, Daniel Heid, Carsten Mueller-Tidow, Max von Kleist, Christoph Dieterich

## Abstract

Nanopore direct RNA sequencing has emerged as a powerful technology for studying RNA and modifications at single-molecule resolution. Although originally developed for polyadenylated RNAs, protocol adaptations have extended its application to non-polyadenylated RNA species, including tRNAs and rRNAs. However, the lack of open and flexible (de)multiplexing solutions remains a limitation for large-scale sequencing studies. Existing community-driven approaches enable multiplexing of polyadenylated RNAs and, more recently, tRNAs, but do not fully address the challenges of targeted sequence-specific Nanopore sequencing of non-polyadenylated RNAs.

Here, we present an improved sequence-specific direct RNA sequencing protocol for ribosomal RNAs, achieving enhanced capture and longer read lengths as demonstrated for human 28S rRNA. We complement this protocol with an expanded WarpDemuX software toolbox for flexible sequence-specific (de)multiplexing. We validate the approach using human cell lines carrying knockouts of different C/D box snoRNAs involved in site-specific 2*′*-O-ribose methylation of 28S rRNA. Our results demonstrate efficient multiplexed detection of sequence-specific RNA molecules and associated modification changes. While established here using rRNA, we validate our approach on another panel of non-polyadenylated RNA species, providing an open and scalable framework for targeted direct RNA sequencing and RNA modification analysis.

## 1 Introduction

Nanopore direct RNA sequencing (DRS) has emerged as a powerful technology for studying RNA modifications at single-molecule resolution. Although DRS was initially developed for the sequencing of polyadenylated RNAs, community driven protocol adaptations also enable the sequencing of non-polyadenylated RNAs such as tRNAs [1, 2, 3, 4] and rRNAs [5, 6]. Despite these advances, the lack of official (de)multiplexing solutions limits the feasibility of large-scale sequencing studies, as well as the broader adoption of the technology. Multiplexing reduces the cost of sequencing by allowing the simultaneous sequencing of multiple samples, which is achieved by incorporating unique identifier sequences (barcodes) into each sample prior to sequencing. For sequencing of polyadenylated RNAs, Oxford Nanopore Technologies (ONT) recently released an official kit. Furthermore, DeePlexiCon [7] is available for the deprecated RNA002 chemistry and two community-developed solutions have been implemented for both, RNA002 and the current RNA004 chemistry: SeqTagger [8] and WarpDemuX [9]. SeqTagger uses a GPU-accelerated RTA basecaller [8], whereas WarpDemuX uses a lightweight Support Vector Machine-Dynamic Time Warping Distance (SVM-DTWD) classifier, which works directly on the raw data [9]. Recently, we expanded this toolkit by an improved and specialized WarpDemuX framework for sequencing of tRNAs [10], which uses Soft-DTW [11] and Barycenter Averaging [12, 13] instead of the standard DTW. This enables the direct classification of barcode fingerprints [10] by a gradient boosted trees classifier. A tRNA-specific SeqTagger model is commercially available. However, both methods do not account for the unique challenges of sequence-specific nanopore sequencing. These are on the one hand the absence of a polyA tract and on the other hand, the presence of a variable DNA-RNA boundary sequence.

The analysis of RNA modifications installed on ribosomal RNAs (rRNAs) is of high interest, as increasing evidence reveals that rRNA modifications are dynamically regulated in development and disease [14, 15, 16]. We have previously presented a sequence-specific approach to sequence native human rRNAs on the nanopore [5, 17]. While capture was highly efficient for the 18S rRNA, 28S rRNA capture and read length remained suboptimal.

Here, we present an improved sequence-specific nanpore rRNA sequencing protocol and an expanded WarpDemuX toolbox for (de)multiplexing of sequence-specific DRS, WarpDemuX-custom. A real-world analysis of human cell lines harboring knockouts for different snoRNAs, required for installation of specific 2’-O ribose methylations on the 28S rRNA, demonstrates the performance of our strategy. Furthermore, we demonstrate the applicability of WarpDemuX-custom to other sequence-specific DRS approaches, such as the sequencing of non-polyadenylated lncRNAs.

## 2 Results

### 2.1 Optimization of direct rRNA sequencing

In our previous work, we designed sequence-specific sequencing adapters (v1) for human rRNAs [5, 17]. While capture of the human 18S rRNA was highly efficient, the 28S rRNA was underrepresented irrespective of the sequencing chemistry used (Supplemental Figure S1A). Read lengths are however markedly increased for both, 18S and 28S rRNA with the state-of-the-art RNA004 chemistry compared to the RNA002 chemistry (Supplemental Figure S1B,C).

The reverse transcription adapter in DRS is ligated to the 3’ end of the RNA with T4 DNA ligase, which can not ligate substrates with gaps or flaps. Sequencing of polyadenylated total RNA revealed that the last 3’ nucleotide annotated in NCBI (e.g. NR_003287.4) is actually not present in human 28S rRNA (Supplemental Figure S1D). Consequently, we designed a new 28S adapter (v2) shifted by one nucleotide, which resulted in a higher fraction of 28S rRNA reads (Figure 1A). Taking the different sequence lengths (28S 5069 nts vs. 18S 1869 nts) and consequently longer sequencing time per molecule into account, the ratio of 28S to 18S is closer to the theoretically expected ratio of 2.7 (assuming that a 18S rRNA molecule occupies the pore 14.4 sec, whereas the 28S rRNA requires 39.0 sec at a sequencing speed of 130 nts/sec). Interestingly, we also observed an improvement of read lengths, for both the 28S and as well the 18S rRNA, evident in eCDF plots (Figure 1C,D), violin plots (Supplemental Figure S1E-F) and IGV coverage tracks (Supplemental Figure S1G-H).

**Figure 1:**
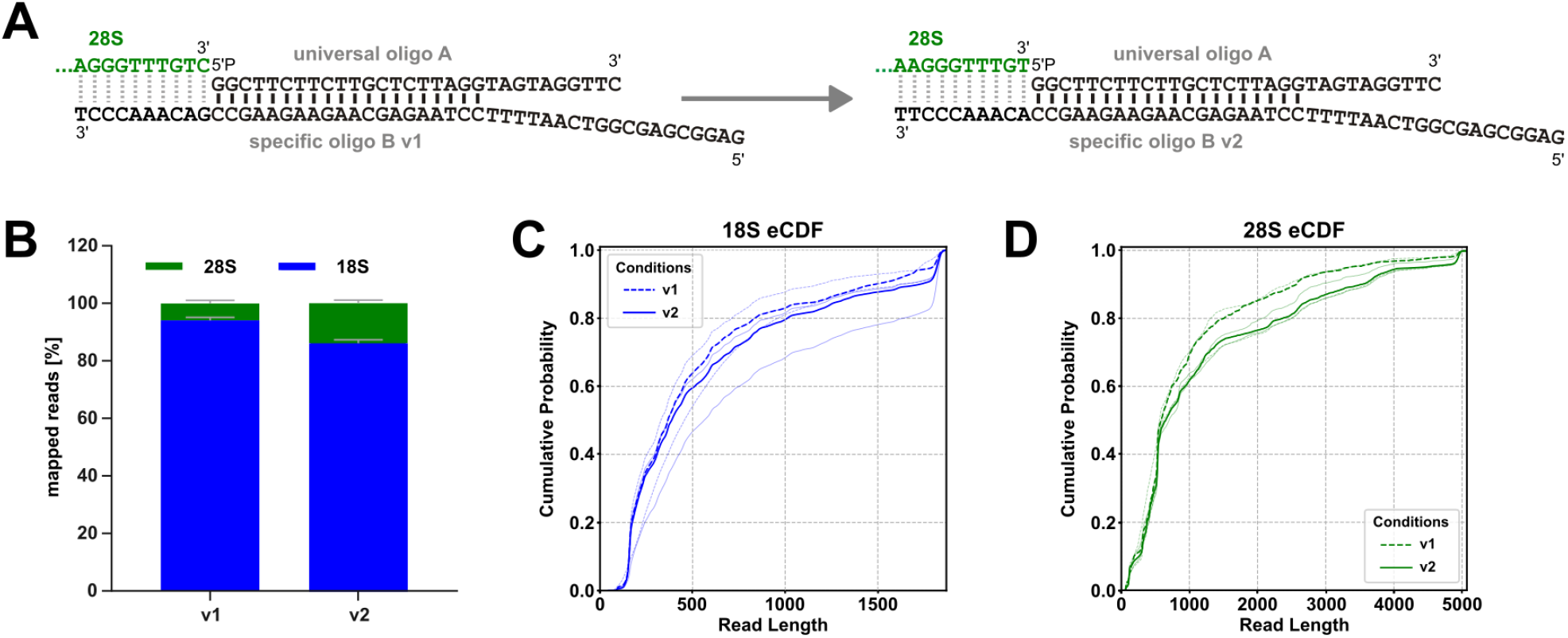
Optimization of direct rRNA-sequencing. **(A)** Optimization of the 28S rRNA custom adapter. Shown is the 3’ end of the 28S rRNA (left = v1, NCBI annotated, right = v2, detected) in green. The sequence-specific reverse transcription adapter (RTA), which is ligated with T4 DNA ligase to the RNA 3’ end is shown in black. Not shown is the sequencing adapter with motor protein (RLA), which is ligated to the 3’ end of the RTA. **(B)** Proportion of mapped reads for 18S and 28S rRNA sequencing with v1 or v2 adapters. Kasumi-1 wildtype cells, n = 3, RNA004 chemistry. **(C)** eCDF of 18S read lengths with v1 and v2 adapters. Shown are the median (blue) and individual runs (light blue). **(D)** eCDF of 28S read lengths with v1 and v2 adapters. Shown are the median (green) and individual runs (light green). Alternative representations of read lengths (IGV coverage tracks and violin plots) are provided in Supplemental Figure S1E-H.

### 2.2 WarpDemuX-custom

WarpDemuX [9] uses the polyA tail and RNA-DNA boundary to identify the barcode end. WarpDemuX-tRNA [10] similarly exploits the three dAs present in the tRNA-sequencing-specific RNA splint adapter as a characteristic adapter-derived signal. In the case of sequence-specific sequencing of non-polyadenylated RNAs, a polyA tract is absent. The only signal that can be used to identify the barcode end is the DNA-RNA boundary, which however will be different for every sequenced RNA and consequently would require training of novel models for every targeted sequence. This lack of a universal adapter-derived signal prevents robust generalization across experiments targeting different non-polyadenylated RNA species.

To train a general model that can be used for de-multiplexing of non-polyadenylated RNA species, we designed a novel DNA-RNA hybrid adapter. This adapter contains a stretch of five rAs in the universal DNA oligo A, which is ligated to the RNA and sequenced. The sequence-specific oligo B contains a complementary dT stretch and is a standard DNA oligo (Supplemental Figure S2A). The rA stretch introduces a constant and readily identifiable terminus signal within the adapter (Figure 2A), buffering the transition between the barcode and the RNA. This minimizes sequence-dependent effects at the junction and makes barcode segmentation independent from transcript-specific sequence composition, thus enabling robust generalization across different RNA sequences.

**Figure 2:**
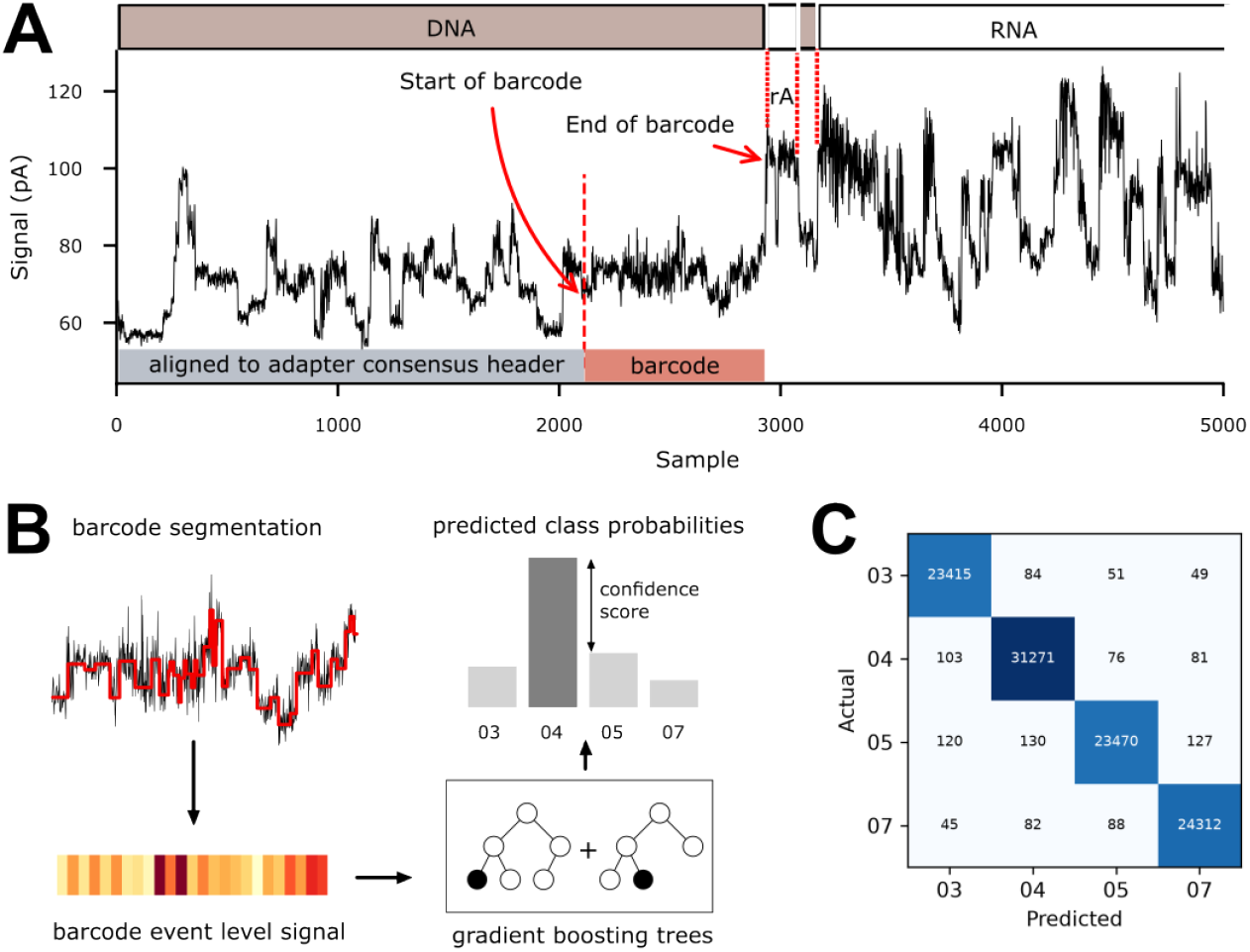
Barcode demultiplexing workflow and performance. **(A)** Raw nanopore signal trace from a single read (first 5000 samples) showing the DNA sequencing adapter and RNA molecule regions, within the DNA adapter the consensus header region (light gray) and barcode region (light red/orange). The start and end of the barcode, the rA stretch and end of the adapter are marked with red dashed vertical lines. **(B)** Barcode processing pipeline: (top-left) event detection in the extracted barcode signal, (bottom-left) segmented signal representation of the barcode region (mean per event), (bottom-right) classification using gradient boosted trees and (top-right) predictive distribution over barcode classes with classification confidence score used for filtering. **(C)** Classification performance on the test set.

This adapter design extends WarpDemuX to non-polyadenylated targets while preserving cost efficiency. The sequence-specific oligo is a standard DNA oligo, whereas the universal hybrid oligo can be reused across sequence contexts, requiring redesign only of the inexpensive sequence-specific DNA component when targeting new RNAs.

We trained WarpDemuX-custom with four barcodes (03, 04, 05, 07 from van der Toorn *et al*. [9]) on two sets of four *in vitro* transcribed RNAs (IVTs), which included but were not restricted to human rRNAs (Supplemental Figure S2B and Methods). Two IVTs were present in both training sets but with shuffled barcodes. The barcode processing pipeline is shown in Figure 2B. The confusion matrix shows efficient de-multiplexing on the test set (Figure 2C, see Methods and Supplemental Figure S2B for details on read splitting. Precision and Recall are high (99%) along with a high read recovery (Table 1).

**Table 1:** Model performance on test set. Precision, recall and recovery are calculated after filtering predictions with confidence scores above the barcode-specific threshold, calibrated on the validation set to reach 99% precision per barcode: 0.080, 0.317, 0.276 and 0.392 for barcodes 03, 04, 05 and 07. Recovery is the percentage of a barcode’s reads retained by that filter. Supplemental Table S3 shows the effect of applying a single threshold instead.

| Barcode | Precision | Recall | Recovery (%) |
| --- | --- | --- | --- |
| 03 | 0.989 | 0.992 | 99.58 |
| 04 | 0.991 | 0.992 | 99.31 |
| 05 | 0.991 | 0.984 | 98.97 |
| 07 | 0.990 | 0.991 | 99.01 |

### 2.3 Multiplexed analysis of SNORD KO cell lines

Finally, we combined the optimized direct rRNA-seq protocol and WarpDemuX-custom and designed barcoded custom adapters for the human 18S and 28S rRNAs. As use case for our new sequencing and (de)multiplexing strategy, we used single-cell derived biallelic knockouts of three different C/D box snoRNAs with known 28S rRNA 2’-O-ribose methylation sites: *SNORD74* installs Cm_3841_, *SNORD76* Am_2363_ and *SNORD81* Am_400_ (28S rRNA nucleotide numbering according to [18] (Figure 3A). We performed multiplexed sequencing of four biologicial samples: wildtype (WT) Kasumi-1 cells and the three different knockouts (KO) of individual snoRNAs. Libraries were split after ligation of the barcoded RTA adapter and samples either sequenced in individual runs (Supplemental Figure S3) to derive the ground truth or subjected to multiplexed sequencing (Figure 3B). Data were basecalled with Dorado v1.1.1 and all available modification models. This analysis clearly revealed that the respective modifications are only absent in the respective KOs, and basecalling statistics in the three other cell lines at the positions of interest are very similar (Figure 3B), indicating efficient de-multiplexing.

**Figure 3:**
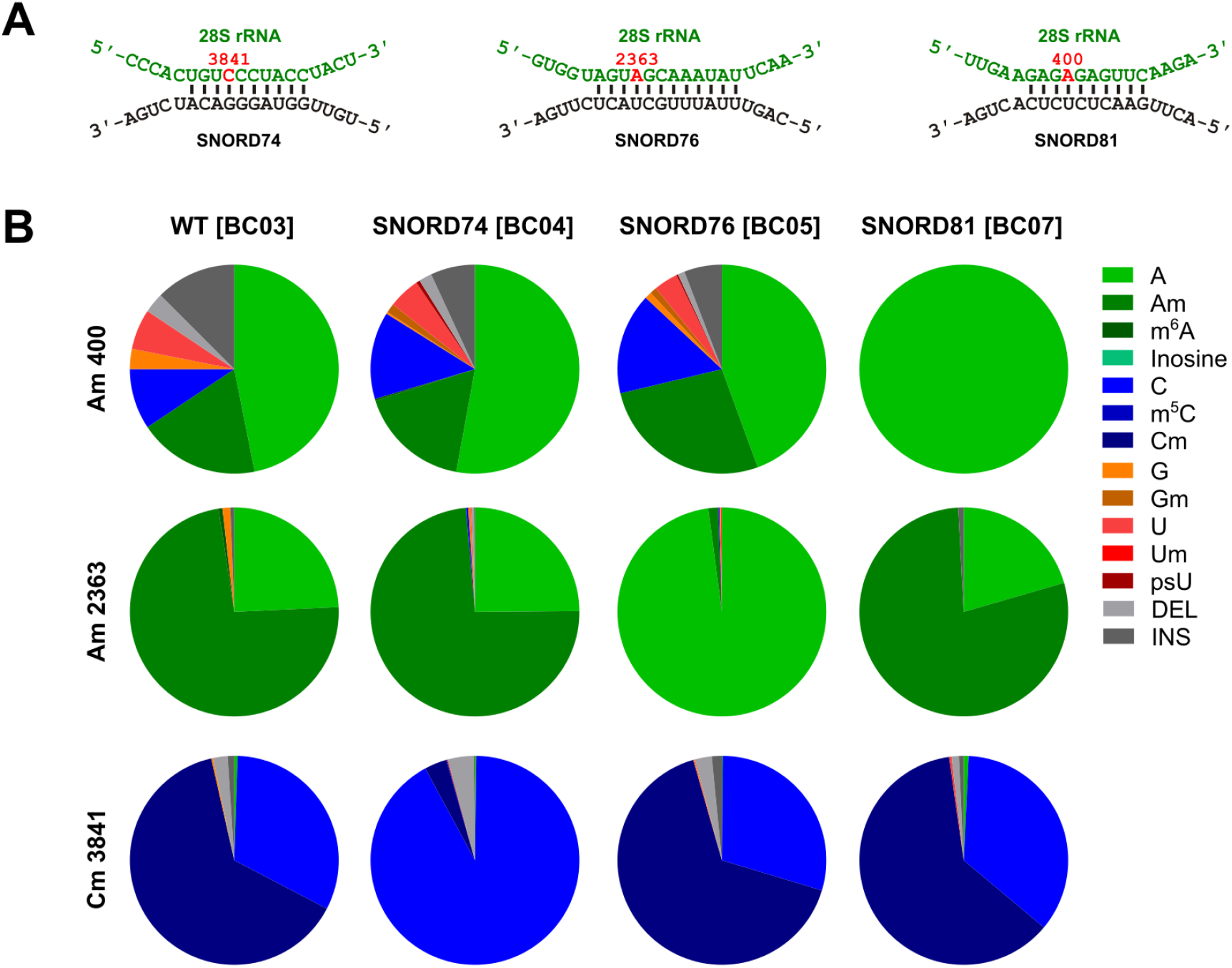
Multiplexed sequencing of Kasumi-1 SNORD KO cell lines. **(A)** 28S rRNA sequences and complementary SNORD RNA sequences. The nucleotide positions correspond to [18] and the snoRNA binding sites were derived from snoRNA-LBME-db [19]. **(B)** Dorado v1.1.1 basecalling statistics at the target positions from multiplexed rRNA sequencing with the indicated barcodes and WDX-custom demultiplexing. The corresponding basecalling statistics of the individual sequencing runs are given in Supplemental Figure S3.

### 2.4 Sequence-specific multiplexing of non-polyadenylated RNAs

WarpDemuX-custom is agnostic to the targeted RNA and can therefor be used to sequence individual non-polyadenylated lncRNAs in a total RNA preparation. To demonstrate the broad applicability, we targeted several non-polyadenylated cellular RNAs with different barcode adapters, inspired by the analysis of [20]. *TERC* is the RNA component of telomerase, *RPPH1* the RNA component of RNase P and *RMRP* is the RNA component of mitochondrial RNA processing endoribonuclease. Furthermore, we have included *RN7SK*, which is part of the 7SK snRNP. For this proof-of-principle analysis, we sequenced these four RNAs with different barcodes in one biological sample, here HeLa total RNA. The four target RNAs were ligated in a multiplexed reaction to the barcodes as indicated (Figure 4A, upper panel). The confusion matrix

**Figure 4:**
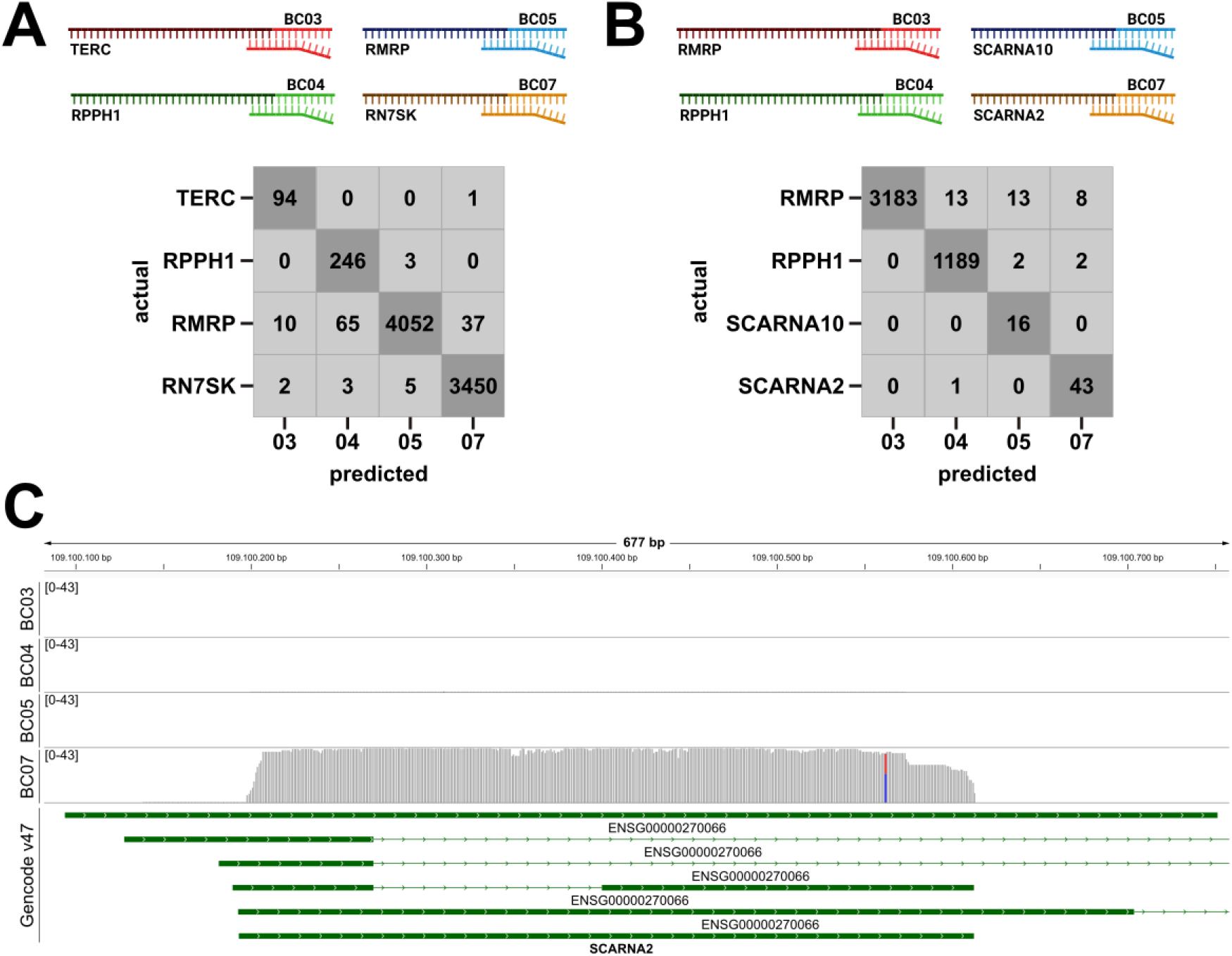
Multiplexed sequencing of non-polyadenylated lncRNAs. **(A)** Upper panel: Experimental design of multiplexed sequencing of non-polyadenylated cellular RNAs. HeLa total RNA was subjected to sequence-specific DRS with four barcoded adapters that target *TERC, RPPH1, RMRP* and *RN7SK* as indicated. Lower panel: confusion matrix of aligned reads mapping to the actual and predicted gene loci for the different barcodes. **(B)** Upper panel: Multiplexed sequencing of *RMRP, RPPH1, SCARNA10* and *SCARNA2* as in A. Lower panel: confusion matrix of aligned reads mapping to the actual and predicted gene loci for the different barcodes. **(C)** IGV coverage track of chr1:109,100,082-109,100,757 covering the *SCARNA2* locus. Annotation and color coding are derived from the Gencode v47 assembly (https://genome.ucsc.edu/cgi-bin/hgTrackUi?hgsid=3643038459_aGxX0KHw4d4A28j16AAMppqRu3lo&db=hg38&g=knownGeneV47). Reads in both confusion matrices were filtered at a confidence score threshold of 0.9.

(Figure 4A, lower panel) shows efficient de-multiplexing. Next, we included two Small-cajal body specific RNAs, *SCARNA2* and *SCARNA10* to demonstrate the performance of WarpDemuX-custom in a highly unbalanced data set (Figure 4B) - *SCARNA2* and *SCARNA10* have very low expression levels, compared to RMRP, which shows some leakage to other barcodes, irrespective of the barcode used to target RMRP. Figure 4C shows the exemplary IGV coverage track of *SCARNA2*, which was targeted with the barcode 07 adapter. This clearly shows that we capture specifically the *SCARNA2* RNA and no RNAs derived from overlapping gene loci. The coverage tracks for the other targeted RNAs show a similiar pattern (Supplemental Figures S4 and S5).

## 3 Discussion

### 3.1 Optimized direct rRNA-seq protocol

In this work, we present an updated and improved direct RNA-sequencing protocol for human ribosomal RNA. Two general strategies for rRNA-DRS exist: enzymatic polyadenylation of total RNA, followed by sequencing with the standard adapter [21, 14] and sequence-specific library preparation with custom adapters [6, 5]. We analyzed the 3’ read ends for both approaches (Supplemental Figure S6) and found for the 18S (see also [5]) and 28S rRNA a substantial fraction of missing 3’ read ends with the polyadenylation approach. To increase the fraction of mature, full length reads, we decided to optimize our sequence-specific approach. A detailed analysis of the polyadenylated dataset revealed interestingly, that the last nucleotide in the 28S rRNA reference is actually absent in human cells (Supplemental Figure S1D). The dominant end position is nt 5069, but we also see a fraction of reads ending at nts 5066-5068. Consequently, our previous adapter targeting nt 5070 could not be efficiently ligated by T4 DNA ligase, which does not tolerate gaps. In order to select for full-length molecules, we decided to use only one adapter targeting nt 5069, but the usage of an adapter mixture would be feasible as well. Due to the shorter pore occupancy time, we expect a bias towards shorter 18S rRNA molecules, which we could reduce with the new 28S rRNA adapter (Figure 1B). In summary, the updated RNA004 sequencing chemistry, reverse transcription with Induro reverse transcriptase and the new 28S rRNA adapter result in a higher 28S rRNA coverage and overall increased read lengths.

### 3.2 WarpDemuX-custom

Direct RNA sequencing is state-of-the-art and the most promising method for current and future research in the area of epitranscriptomics. The technology lacks however important tools, such as algorithms to detect a wide variety of RNA modifications and official barcoding options for non-polyadenylated RNAs. An official barcoding kit for poly(A) sequencing was released by ONT recently. Furthermore, community driven solutions for sequencing of polyadenylated RNAs (i.e. mRNAs) and tRNAs have been presented by two labs in the past (namely SeqTagger [8] and WarpDemuX [9, 10]), whereas no solution exists for sequence-specific DRS. However, the option of multiplexed sequencing is of particular importance for use cases like direct rRNA sequencing, and even more for less abundant non-polyadenylated ncRNAs. Here, the overall sequence complexity is low, which would enable the pooled sequencing of several samples on one flow cell.

In this work, we present WarpDemuX-custom, which extends the WarpDemuX toolkit and allows multiplexed sequencing of all RNA types that are accessible to sequence-specific DRS. We introduce a short RNA stretch into the barcoded DNA adapter, which is used to identify the barcode signal in the signal trace (Figure 2). The use of this constant signal renders the WarpDemuX-custom model agnostic to the RNA that is sequenced. In other words, the same model can be used for example to de-multiplex rRNA runs from different species. Furthermore, also other non-polyadenylated RNA species can be targeted, even in one sample (see Figure 4). We demonstrate that WarpDemuX-custom works over a wide range of expression levels. Only at very high expression levels (*RMRP*) we detected a small number of mis-assigned reads with both tested barcodes (Figure 4A,B). A real world example, using the latest Dorado basecalling models, demonstrates the performance of WarpDemuX-custom for rRNA (Figure 3). In the current version, WarpDemuX-custom supports four barcodes, but it could be extended to up to 10 barcodes, which have been described for the original WarpDemuX [9].

In summary, we present here for the first time an open-source solution for the multiplexed sequencing of non-polyadenylated RNAs, WarpDemuX-custom along with an optimized protocol for sequence-specific sequencing of human rRNAs. We have demonstrated the performance of WarpDemuX-custom for rRNAs and non-polyadenylated cellular lncRNAs, but due to the sequence-agnostic design it is applicable to all non-polyadenylated RNA species with defined 3’ ends, including gRNA from non-polyadenylated RNA viruses [22]. Multiplexed, targeted sequencing reduces the sequencing cost, enables a higher throughput, and can also increase the output per flow cell due to a higher pore occupancy. In terms of costs, only specific adapter oligos are required in addition to the standard reagents. The slightly more expensive barcoded adapter with the rA stretch is the universal oligo, whereas the sequence-specific oligos are low-cost standard DNA oligos. We have not tested different input amounts systematically here, but based on our experience lower input amounts per sample are expected to be sufficient for multiplexed rRNA sequencing, which is especially relevant for clinical samples. WarpDemuX-custom is available at https://github.com/dieterich-lab/WarpDemuX.

## 4 Material and Methods

### 4.1 Cell culture and generation of KO cell lines

Plasmids encoding sgRNAs upstream and downstream of the respective snoRNA locus were generated by cloning the corresponding sequence into the pL-CRISPR.EFS.GFP vector (Addgene). Plasmids encoding sgRNAs targeting sequences upstream and downstream of the respective target were electroporated (Neon Transfection System, Thermo Fisher Scientific) at equal amounts into Kasumi-1 cells. 24-48h after electroporation GFP-positive cells were FACS-sorted as single cells into 96-well dishes. Grown single-cell derived clones were screened for biallelic target knockout on the DNA level by flanking PCRs and on the RNA level by RNA extraction and RT-qPCR.

HeLa cells were obtained from the DSMZ-German Collection of Microorganisms and Cell Cultures GmbH and cultured in D-MEM (Thermo Fisher Scientific) supplemented with 10% fetal bovine serume (Merck) and 1x penicillin/ streptomycin (Thermo Fisher Scientific) at 37°C and 5% CO_2_. Total RNA was isolated with TRIzol (Thermo Fisher Scientific) according to the manufacturers protocol. Nanodrop (Thermo Fisher Scientific) and Fragment Analyzer (Agilent Technologies) were used to assess quantity and quality of the isolated RNA.

### 4.2 Generation of *in vitro* transcripts

*In vitro* transcripts (IVTs) were generated from PCR products containing a T7 promoter sequence. The 18S IVT [5], 28S-F3 IVT [17] and blaR IVT [23] have been described previously. T7-ampR was amplified from pBSIIKS, T7-lacI from pET16b and T7-EGFP from pEGFP-C1 with primers listed in Supplemental Table S4, with the respective forward primers encoding a T7 promoter sequence as described in [23]. The MegaScript T7 transcription kit was used to generate IVTs from the PCR templates. IVT reactions were cleaned up using RNA Clean & Concentrator kit-100 (Zymo Research).

### 4.3 Design of sequence-specific barcoded adapters

Barcode sequences are derived from [9]. We have trained here a model for four barcode sequences (03, 04, 05, 07). The sequences of the barcoded adapters was extended by a stretch of five rA’s (see Supplemental Figure S2), which was placed directly adjacent to the barcode sequence. To ensure efficient ligation by T4 DNA ligase, the terminal 5’ nucleotides are DNA nucleotides.

The sequence for targeting the 18S rRNA has been described previously [5] and is based on the 18S rRNA sequence deposited in NCBI (NR_003286.4). The 28S rRNA adapter was designed based on the NCBI sequence (NR_00328.4) and the sequencing of polyadenylated 28S rRNA (see 4.1). The *TERC, SCARNA2* and *SCARNA10* sequences were obtained from ENSEMBL. The exact 3’prime end of *RMRP* was derived from [24] and that of *RPPH1* from [25]. The adapter sequence for *RN7SK* was obtained from [26]. For the IVTs used for the training datasets, the 3’end sequences were obtained from the reverse PCR primer sequences (see Supplemental Table S4).

### 4.4 Nanopore direct RNA-sequencing

To generate the training and test data sets (IVTs and Kasumi-1 rRNA), one library was split for individual and multiplex runs. First, 200 ng RNA were ligated to the respective WDXC adapter using T4 DNA ligase (New England Biolabs) as described previously [5]. Reverse transcription (to dissolve secondary structures) was done using Induro reverse transcriptase (New England Biolabs). After reverse transcription, the reaction was quenched by addition of EDTA to a final concentration of 25 mM EDTA. Reactions were cleaned up using 1.8x RNA Clean XP beads (Beckman Coulter). Before ligation of the RLA adapter (SQK-RNA004, Oxford Nanopore Technologies), the samples were split: 75% were used for the individual runs and 25% were pooled for the multiplex runs.

For the sequencing of the non-polyadenylated RNAs, 200 ng total RNA was subjected to T4 DNA ligation with a mixture of all four barcoded adapters. Further library preparation was performed as described above without splitting of the library.

After the final clean up with RNA Clean XP beads, the libraries were loaded on FLO-PRO004RA flow cells (Oxford Nanopore Technologies) and sequenced to a data target of 50 Mb estimated bases, with high accuracy basecalling enabled on a Promethion P24-A100 device. For comparison of the two 28S adapters, split libraries were sequenced on different devices (P24, P2 Solo, MinION). For analysis of RNA modification, data were re-basecalled with Dorado v1.1.1 and v5.2 RNA models. Data were aligned to the respective reference sequences with minimap2. Integrative Genomics Viewer was used for Coverage Track visualization and custom Phython scripts for eCDF and violin plots.

Universal barcoded oligos A of the sequencing adapter are listed in Supplemental Table S5. Sequence-specific barcoded oligos B of the sequencing adapter are given in Supplemental Table S6. Adapters were annealed as described previously [5].

### 4.5 WarpDemuX-custom training

#### 4.5.1 Data splitting

Our analysis combines two independent training datasets, which contained rRNA but also non-related IVT sequences (Supplemental Figure S2). Composition of set 1: 18S IVT - BC03, 28S-F3 IVT - BC04, lacI - BC05, ampR - BC07. Composition of set 2: EGFP - BC03, blaR - BC04, 18S IVT - BC05, lacI - BC07. The number of mapped reads per dataset are given in Supplemental Table S1. For each replicate, reads were split into training, calibration/validation, and test sets according to the ratio 4:1:2 (training:calibration/validation:test) using a stratified approach. After splitting, replicates were combined within each split to form the final training, calibration/validation, and test sets. Mapped reads per label and data split are given in Supplemental Table S2.

#### 4.5.2 Barcode fingerprint generation

Barcode fingerprint generation follows the same methodology as described in [10], with parameters tuned on the training set and distributed with the model as its signal processing configuration (rna004_130bps@v1.0_custom, WarpDemuX v1.3.0).

#### 4.5.3 Barcode classification

We use gradient boosting trees (as implemented in CatBoost [27]) for barcode classification. Training (*<*1minute) was performed on the refined barcode fingerprints with early stopping on the calibration/validation set (100 rounds without improvement), using balanced class weights, a learning rate of 0.1, a maximum tree depth of 6 and at least 5 samples per leaf and a fixed random seed of 42. The best model based on validation performance was selected for evaluation on the test set (`WDX4_custom_rna004_v1_1`).

#### 4.5.4 Confidence score calibration

For each prediction, we compute a confidence score as the difference between the highest and second-highest class probabilities [9]. We calibrate barcode-specific confidence thresholds on the held-out calibration/validation set to achieve 99% precision for each barcode class. As in [9, 10], predictions with confidence scores below these class-specific thresholds are filtered out as “unclassified”. The model performance is then evaluated on the independent test set. The calibrated thresholds are 0.080, 0.317, 0.276 and 0.392 for barcodes 03, 04, 05 and 07 respectively. For the cellular RNA data in Figures 3 and 4 we instead applied a single confidence score threshold of 0.9 to all four barcodes, which is stricter than every per-barcode calibrated threshold; Supplemental Table S3 shows the effect of various thresholds on the test set.

## 5 Acknowledgments

We are grateful to Harald Wilhelmi for his excellent assistance in supporting software development and running and maintaining the Dieterich Lab compute cluster.

## 6 Supplementary data

Supplementary data is available online.

## 7 Conflicts of interest

The authors declare that they have no competing interests.

## 8 Funding

This work was funded by a DFG grant to CD (TRR319 RMaP project no. 439669440). CD acknowledges additional funding by the Klaus Tschira Stiftung gGmbH (grant #00.013.2021). MFB and CMT acknowledge funding by the German Research Foundation project number 533056198 SFB 1709.

## 9 Data availability

All Nanopore sequencing data and software freeze (v1.3.0) are available on Zenodo under https://doi.org/10.5281/zenodo.18983222. The latest WarpDemuX-custom models can be found at: https://github.com/dieterich-lab/WarpDemuX.

## 10 Author contributions statement

INV and WT developed the WarpDemuX-custom barcode design. INV performed all experiments unless otherwise stated. WT and SM trained WarpDemuX-custom. SM analyzed data. MFB and DH generated the Kasumi-1 SNORD KO cell lines and isolated RNA with supervision of CMT. MK and CD designed the study. INV and WT wrote the manuscript, with input from all authors.

## Notes

### Competing Interest Statement

The authors have declared no competing interest.

